# Shavenbaby protein isoforms orchestrate the self-renewal *versus* differentiation of *Drosophila* intestinal stem cells

**DOI:** 10.1101/627554

**Authors:** Sandy Al Hayek, Ahmad Alsawadi, Zakaria Kambris, Jean-Philippe Boquete, Jérôme Bohère, Brice Ronsin, Serge Plaza, Bruno Lemaitre, François Payre, Dani Osman

**Affiliations:** Faculty of Sciences III, Lebanese University, Tripoli, 1300, Lebanon; Azm Center for Research in Biotechnology and its Applications, LBA3B, EDST, Lebanese University, Tripoli, 1300, Lebanon; Centre de Biologie du Développement (CBD), Centre de Biologie Intégrative (CBI), Université de Toulouse, CNRS, Bat 4R3, 118 route de Narbonne, F-31062, Toulouse, France; Biology Department, Faculty of Arts and Sciences, American University of Beirut, Beirut, Lebanon; Global Health Institute, School of Life Sciences, Station 19, EPFL, 1015 Lausanne, Switzerland

## Abstract

Signaling pathways are key regulators of adult stem cell homeostasis and underlying mechanisms are often deregulated in cancers. Recent studies of epithelial tumors have involved OvoL/Svb transcription factors, which produce isoforms with antagonistic activities. Here we show that Svb, the unique OvoL factor in *Drosophila*, directly integrates multiple signaling inputs to coordinate the behavior of adult intestinal stem cell lineage. Under steady state, Svb mediates Wnt and EGFR signaling to ensure stem cell renewal and progenitor survival. This requires the post-translational processing of Svb into a transcriptional activator by Polished rice (Pri) regulatory peptides, under the regulation of ecdysone signaling. In response to PDM1, Svb expression is specifically maintained in enterocytes where it acts as a transcriptional repressor sufficient to override mitogenic signals and impose differentiation. Altogether, these results demonstrate that the OvoL/Svb transcriptional switch controls the balance between stem cell survival, self-renewal and differentiation.

## INTRODUCTION

Living organisms are constantly exposed to internal and environmental challenges that may disturb specific cell functions and promote cell death. To maintain homeostasis, most organs are regenerated by stem cells that self-renew and differentiate to replenish damaged tissues by replacing dead cells. The highly regenerative digestive system is kept intact during adulthood by the activity of intestinal stem cells residing in the gut epithelium. Recent findings on intestinal stem cells in flies have greatly advanced our understanding of the regulatory signaling networks underlying stem cell biology and their implication in cancers (reviewed in (Herrera and Bach, 2019; Li and Jasper, 2016; Perochon et al., 2018)).

The adult *Drosophila* midgut consists of a highly compartmentalized epithelium (Buchon et al., 2013), which shares anatomical and physiological similarities with the mammalian counterpart (Casali and Batlle, 2009). The fly intestinal stem cells (ISCs) are scattered underneath the basal surface of the midgut epithelium (Micchelli and Perrimon, 2006; Ohlstein and Spradling, 2007). In most cases, ISC divide asymmetrically to generate a new stem cell and a progenitor called enteroblast (EB) (de Navascues et al., 2012; Goulas et al., 2012; Ohlstein and Spradling, 2007). EB are post-mitotic cells that progressively acquire characteristics of either absorptive enterocytes (EC) or hormone-secreting enteroendocrine cells (EE) (Ohlstein and Spradling, 2007). It has been proposed that EE arise from a separate pool of progenitors, called pre-enteroendocrines, which express markers of both ISCs and EEs (Biteau et al., 2011; Zeng and Hou, 2015).

The evolutionarily conserved Notch pathway establishes the asymmetry between ISC and EB after division (Ohlstein and Spradling, 2007; Perdigoto et al., 2011). ISCs express Delta, a transmembrane ligand that activates the Notch receptor in neighboring daughter EBs, with high levels of Notch promoting the EC fate, whereas EE requires lower levels of Notch (Ohlstein and Spradling, 2007). The JAK/STAT pathway acts downstream Notch to ensure differentiation of committed progenitors (Beebe et al., 2010; Jiang et al., 2009).

Gut homeostasis further relies on a tight regulation of ISC division through cooperative activity of additional signaling pathways (reviewed in (Buchon and Osman, 2015)). For instance, the epidermal growth factor receptor (EGFR), Wingless (Wg) and JAK-STAT pathways contribute to ISC preservation and proliferation in a redundant manner, and loss of all three pathways is needed to abrogate ISC maintenance (Biteau and Jasper, 2011; Jiang and Edgar, 2009; Jiang et al., 2011). Despite the wealth of knowledge accumulated on the role of signaling pathways in regulating ISC maintenance, division and differentiation, the intrinsic mechanisms by which ISCs integrate these cues remain poorly understood.

Previous work has shown that Wnt and EGFR pathways specify embryonic epidermis differentiation through controlling the expression of Ovo/Shavenbaby (Svb), a transcription factor (Payre et al., 1999) that governs cell remodeling. In response to small peptides called Polished rice (Pri), encoded by an atypical RNA that contains only four small open reading frames (smORF), Svb switches from a long transcriptional repressor (Svb^REP^) to a short activator (Svb^ACT^), following post-translational processing (Kondo et al., 2010; Zanet et al., 2015). Pri expression is directly activated by ecdysone (Chanut-Delalande et al., 2014), the main steroid hormone in insects, which regulates reproduction, sleep, nutritional state and stress resistance (Uryu et al., 2015).

Initially isolated in flies for its role in epidermal differentiation and germ cell maintenance, the Ovo/Svb gene was soon after identified in mammals (Dai et al., 1998) in which three paralogs are called OvoL1-3. OvoL/Svb are specific to animals and encode transcription factors with a conserved C-terminal DNA binding domain, and varying N-terminal regions conferring antagonistic transcriptional activities (Kumar et al., 2012). Human tumor profiling has recently identified OvoL/Svb as regulators of the metastatic potential of many epithelial cancers. Besides pathological situations, functional studies have disclosed important functions of OvoL/Svb for the repair of epithelial tissues, *e.g.* for mammary (Watanabe et al., 2014) and epidermal (Haensel et al., 2019) regeneration in mammals. OvoL/Svb factors are also critical for maintenance of human corneal epithelial cells and in eye regeneration in invertebrates (Kitazawa et al., 2016; Lapan and Reddien, 2012). Moreover, we recently found that Svb is required for the survival of renal nephric stem cells (RNSC) in adult Malpighian tubules. Svb activates the expression of the *Drosophila* anti apoptotic protein (DIAP1) in RNSC by direct interaction with Yorkie, a nuclear effector of the Hippo pathway (Bohere et al., 2018).

In this study, we investigated whether Svb plays a role in sustaining adult *Drosophila* midgut homeostasis in order to regulate ISC survival, division and/or differentiation. We found that *svb* is expressed in both ISC/EBs and ECs, where its inactivation induces apoptotic cell death. Svb directly acts downstream of EGFR and Wnt mitogenic signaling in progenitors, where it is processed into Svb^ACT^ through the activity of Pri/Ubr3/proteasomal pathway. In contrast, Svb^REP^ promotes differentiation of ISC/EB lineage toward ECs, where it is required to maintain functional organization of mature cells. Importantly, Svb^REP^ is a potent inhibitor of tumor induced by deregulated Notch and JAK/STAT pathways, imposing differentiation towards the enterocyte fate. Thus, these findings show that, though integrating multiple signaling pathways and transcription factors, Svb coordinates ISC survival, renewal and differentiation.

## RESULTS

### Svb is required to maintain *Drosophila* adult midgut progenitors

To monitor *svb* expression in the adult midgut, we tested the main enhancers that drive *svb* transcription in the embryo (Frankel et al., 2011). We found that the medial *svb* enhancer drives specific expression in *escargot*-positive cells, therefore identifying them as stem cells and enteroblasts (Figures 1A,B and S1A). Further dissection restricted the cis-regulatory region responsible for progenitor expression to two separate enhancers called *E3-14* (292 bp) and *E6* (1kb).

As a first step to investigate the function of Svb in intestinal stem cells, we used targeted RNAi-mediated depletion at the adult stage. The conditional thermosensitive driver (*esg-Gal4, UAS-GFP, tub-Gal80^ts^*, henceforth referred to as *esg^ts^*) allowed *svb* knockdown in *esg+* cells, resulting in a strong decrease in the number of progenitors (Figure 1C,C’). Using *delta-lacZ* and *Su(H)-GBE-lacZ* markers to discriminate ISCs *vs* EBs respectively, we observed a strong reduction in both stem cells and enteroblasts upon *svb* knockdown (Figure 1C-E’). Similar results were obtained following *svb* depletion specifically either in ISCs (*esg-Gal4, UAS-YFP, Su(H)GBE-Gal80, tub-Gal80^ts^* (Wang et al., 2014), hereafter referred to as *ISC^ts^*) (Figure 1D,D’) or in EBs (*Su(H)-Gal4, tub-Gal80^ts^, UAS-GFP*, hereafter referred to as *Su(H)^ts^*) (Figure S1B). Hence, *svb* depletion causes the loss of both ISCs and EB populations, without detectable effects on enteroendocrine cells (Figure S1B-C).

**Figure 1:**
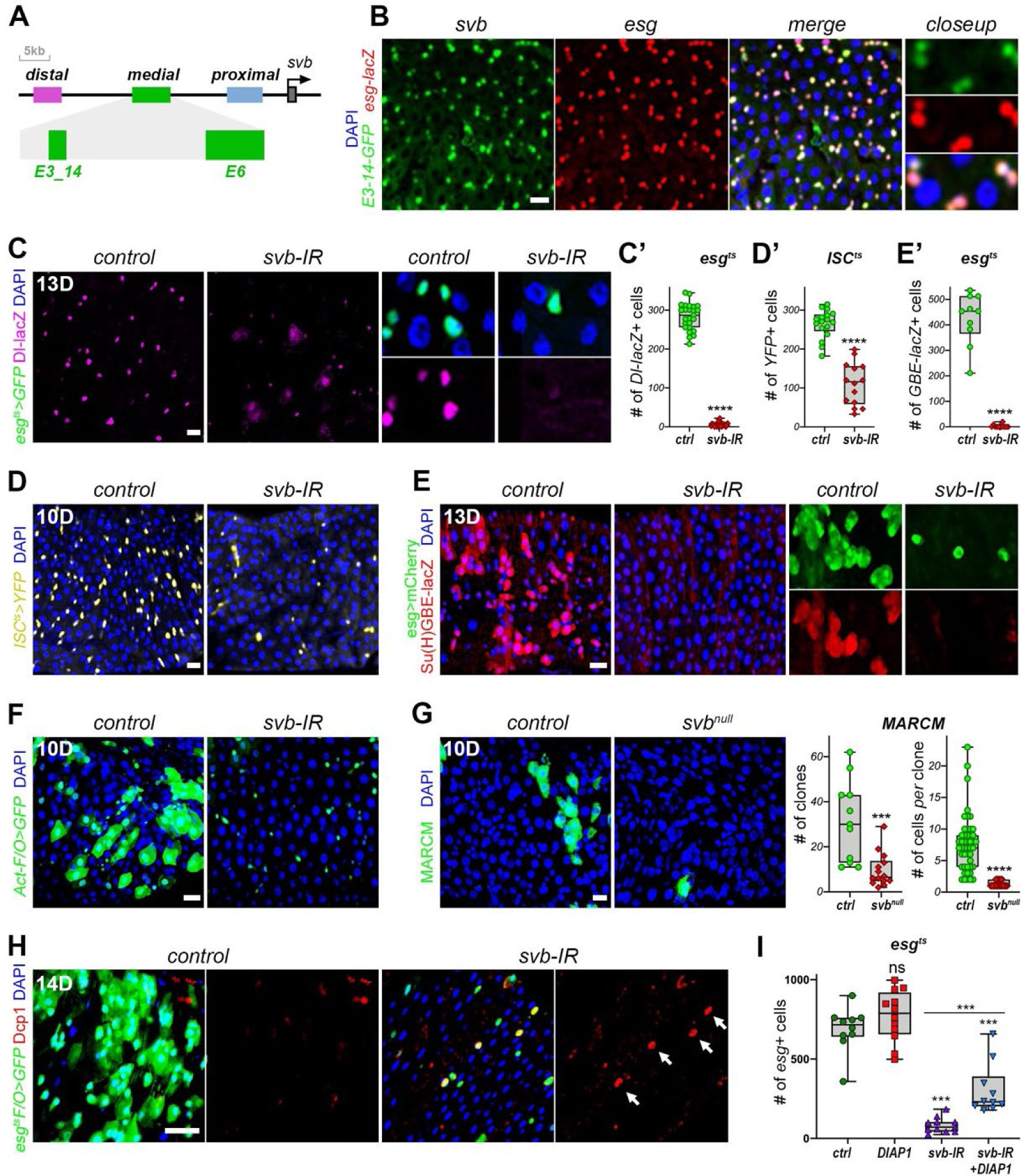
*svb* is expressed in ISC/EBs and is required for their maintenance. (A) Schematic representation of the *svb* locus, with the position of main embryonic enhancers. (B) Expression of GFP (green) driven by the *E3-14 svb* enhancer in ISC/EBs, as shown by co-staining with *esg-lacZ* (β-Gal, red). (C) Expression of *Dl-lacZ* reporter (purple) in *esg^ts^* (control), and *esg^ts^*>*svb*-RNAi. Close ups show separate channels for GFP (green) and Dl-lacZ (purple). (C’) Quantification of *Dl-lacZ*-positive cells in *esg^ts^ svb-*RNAi and control (*esg^ts^>GFP*). (D-D’) Effect of *svb*-RNAi knockdown using *ISC^ts^* (D) and quantification of YFP-positive cells (D’). (E-E’) *Esg^ts^*>*svb*-RNAi in enteroblasts marked by *Su(H)-LacZ* (red), green is *esg^ts^*>*mCherry*; quantification of Su(H)-positive EBs. (F) Actin Flip-out clones of control and *svb*-RNAi expressing cells. Clones are stained by GFP. (G) MARCM clones of control or *svb^R9^* null mutant cells, labeled by GFP (green). Flies were heat shocked once at 37°C for 1 hour and then shifted to 25°C for 10 days. Quantification of the number of clones *per* posterior midgut and number of cells *per* clone. (H) Staining for the apoptotic marker DCP1 (red) in control and *svb*-RNAi *esg-Flo* clones (GFP positive cells, green). (I) Number of GFP-positive cells *per* posterior midgut in *esg^ts^*>GFP (control) and together with DIAP1, *svb-RNAi*, *svb-RNAi* and *DIAP1*. In all panels, DAPI is blue, scale bars represent 20µm. See also Figure S1. In this and all following figures, pictures display representative images of the posterior midgut of female flies aged from 5 to 7 days and shifted to non-permissive temperature for the indicated number of days. P values were estimated by nonparametric Mann-Whitney tests, and Kruskal-Wallis tests with Dunn’s correction, for comparison of two or several samples, respectively. ns: p>0.05, ***: p<0.001, ****: p < 0.0001.

This conclusion was further tested by lineage-tracing using *act^ts^*F/O system (*actin-Gal4; tubGal80^ts^ Act>Cd2> Gal4 UAS-flp UAS-GFP*), which led to random inactivation of *svb* in dividing intestinal cells and their progeny. This experiment showed that *svb* depletion leads to a strong decrease in both the number and size of GFP-labelled clones, when compared to controls (Figure 1F). This was also confirmed by the generation of genetically mutant cells for *svb*, using the MARCM technique (Lee and Luo, 2001). In contrast to controls, we obtained very few clones of cells homozygous for the null *svb^R9^* mutant allele (Delon et al., 2003). In addition, *svb* mutant clones were unable to grow and often restricted to single cells (Figure 1G).

These results therefore suggested that intestinal progenitors lacking *svb* undergo apoptosis, as recently shown for renal stem cells (Bohere et al., 2018). To test this hypothesis, we stained for the apoptotic marker *Drosophila* caspase protein 1 (Dcp1) in intestines bearing clones lacking *svb*, using *esg^ts^F/O* (*esg-Gal4; tubGal80^ts^ Act>Cd2> Gal4 UAS-flp UAS-GFP*) (Jiang et al., 2009) to drive *svb-*RNAi. While only rare Dcp1–positive cells were seen in controls, we observed many apoptosis figures in GFP+ cells upon *svb* depletion (Figure 1H). In addition, whereas overexpression of *Drosophila Inhibitor of Apoptosis-1 (DIAP1)* did not significantly influence the number of *esg+* cells, it partially rescued the loss of ISC/EB when concomitantly expressed with *svb-*RNAi (Figures 1I). Finally, ReDDM lineage tracing (Antonello et al., 2015) further indicated that the loss of stem cells upon *svb* depletion does not results from premature differentiation (Figure S1D).

Taken together, these data show that Svb function is required to protect intestinal stem cells and enteroblasts from apoptosis.

### Pri/Ubr3/proteasomal pathway regulates Svb function in midgut progenitors

Svb is translated as a long (1345aa) repressor (Svb^REP^) that is post-translationally processed into a shorter (946aa) transcriptional activator (Svb^ACT^) (Kondo et al., 2010). This switch is gated by Pri smORF peptides that bind to and activate the E3 ubiquitin ligase Ubr3, triggering its binding to Svb (Zanet et al., 2015). Ubr3 then induces the processing of Svb *via* limited proteasome degradation of its N-terminal repression domain (see Figure 2C). Originally described in epidermal derivatives (Chanut-Delalande et al., 2014; Kondo et al., 2010), a growing body of evidence suggests that Pri-dependent processing underlies Svb function in somatic tissues (Pueyo and Couso, 2011; Ray et al., 2019), including adult stem cells (Bohere et al., 2018).

**Figure 2:**
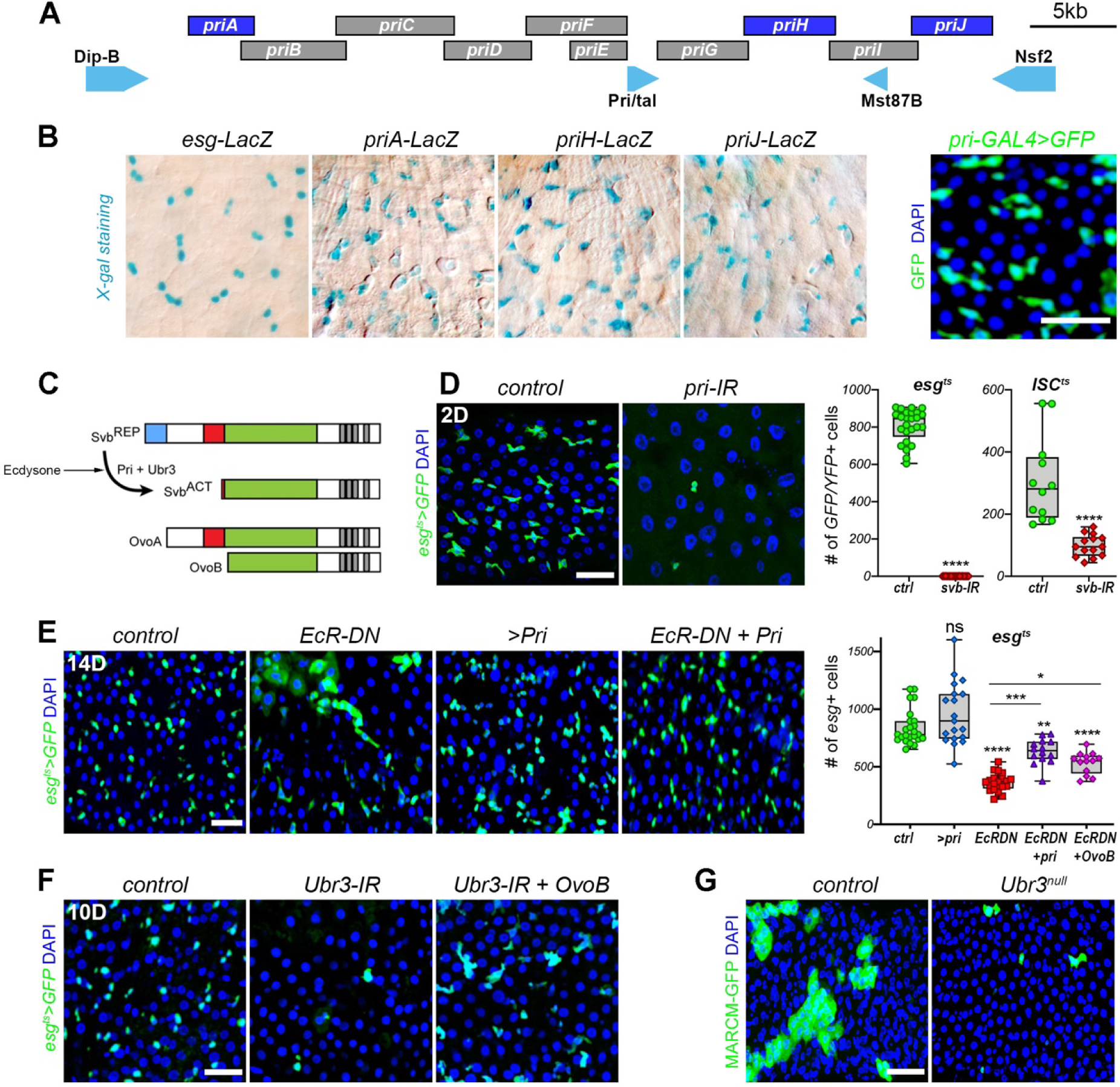
The Pri/proteasome processing of Svb is required for ISC/EB maintenance. (A) Schematic representation of *pri* genomic locus, with the position of Gal4 gene trap insertion. (B) Expression of *priA*, *priJ* or *priH* enhancers (cyan) and *pri-Gal4* transgene (GFP, green). (C) Schematic representation of Svb maturation by proteasomal processing. The germinal isoforms OvoA and OvoB are also shown. (D) Effect of *pri*-RNAi depletion driven by *esg^ts^* and quantification of GFP positive cells *per* posterior midgut. (E) Effect of *esg^ts^* driven expression of *EcRDN*, *Pri*, *EcRDN plus Pri, or EcRDN plus OvoB* and quantification of GFP-positive cells. (F) *Esg^ts^* expression of *Ubr3*-RNAi alone or in combination with *OvoB*. (G) Control and *Ubr3* null MARCM clones, visualized by GFP (green). In all panels, blue is DAPI and scale bars represent 20µm. P values from Mann-Whitney tests (D’) and Kruskal-Wallis tests (E’) are ns:>0.5, *: <0.05, ***: < 0.001, ****: <0.0001. See also Figure S2.

To investigate whether Pri-mediated maturation of Svb was also involved in maintaining ISC/EB, we examined *pri* expression in the adult midgut. Profiling of reporter lines (Chanut-Delalande et al., 2014) showed that three *pri* enhancers (*priA, priJ* and *priH*) drive specific expression in ISC/EBs (Figure 2A,B)*. pri* expression domain was also confirmed by the activity of a gene trap (Galindo et al., 2007) in ISC/EBs (Figure 2B). We next assayed the consequences of *pri* knockdown in ISC/EBs and observed an acute loss of progenitors when Pri-RNAi was driven by *esg^ts^* (Figure 2D), or stem cells when using the *ISC^ts^* driver (Figure S2A). In the epidermis, ecdysone signaling times *pri* expression across development (Chanut-Delalande et al., 2014). We reasoned that if this hormonal control of *pri* expression was occurring in the midgut, cell autonomous disruption of ecdysone signaling should affect the behavior of ISCs. Consistently with this prediction, downregulation of ecdysone signaling *via* a dominant negative receptor (EcR-DN) (Figures 2E and S2D), or RNAi-mediated depletion of EcR (Figure S2B,C), led to a loss of intestinal progenitors. Furthermore, restoring *pri* expression was sufficient to rescue the loss of intestinal progenitors upon EcR inactivation (Figures 2E and S2D). These data thus support that *pri* is an important target of ecdysone required (and somehow sufficient) for the homeostasis of intestinal stem cells.

A key role for Pri in ISC/EBs in triggering the Ubr3-mediated processing of Svb was confirmed by several lines of evidence. First, knockdown of *Ubr3* in intestinal progenitors or stem cells (*ubr3-*RNAi driven by *esg^ts^* or *ISC^ts^*) resulted in a strong reduction of their number (Figures 2F and S2E). Second, similar results were observed in clones of *ubr3* mutant cells, which were very rare and unable to expand (Figure 2G). Third, the loss of Ubr3 could be compensated by expression of the constitutive activator OvoB (Figures 2F and S2E), indicating that Pri and Ubr3 are required in ISCs to trigger the switch of Svb transcriptional activity.

We interpret these results to imply that Svb undergoes Pri/Ubr3-dependent processing in intestinal stem cells to maintain the population of midgut progenitors.

### The Svb activator form triggers ISC proliferation

Having shown that the Svb activator is required for the survival of ISC/EBs, we next investigated if it could be sufficient to influence adult stem cell homeostasis.

Hence, we expressed the OvoB constitutive activator using *esg^ts^* and monitored the behavior of ISCs marked with *delta-LacZ*. OvoB induced a strong increase in the number of ISCs (*esg+, delta-LacZ+* cells), reaching up to four-fold the normal population (Figure 3A). This was due to ISC over-proliferation, as seen by staining for the mitotic marker phosphorylated-histone3 (PH3) (Figure 3A’). Similar results were obtained when OvoB expression was specifically targeted in stem cells using the *ISC^ts^* driver (Figure 3B). Furthermore, if *pri* overexpression has weak if any effects, the simultaneous expression of *pri* and *svb* (to induce the production of Svb^ACT^) provoked a massive proliferation of ISCs, reminiscent of the OvoB phenotype (Figures 3B and S3A).

**Figure 3:**
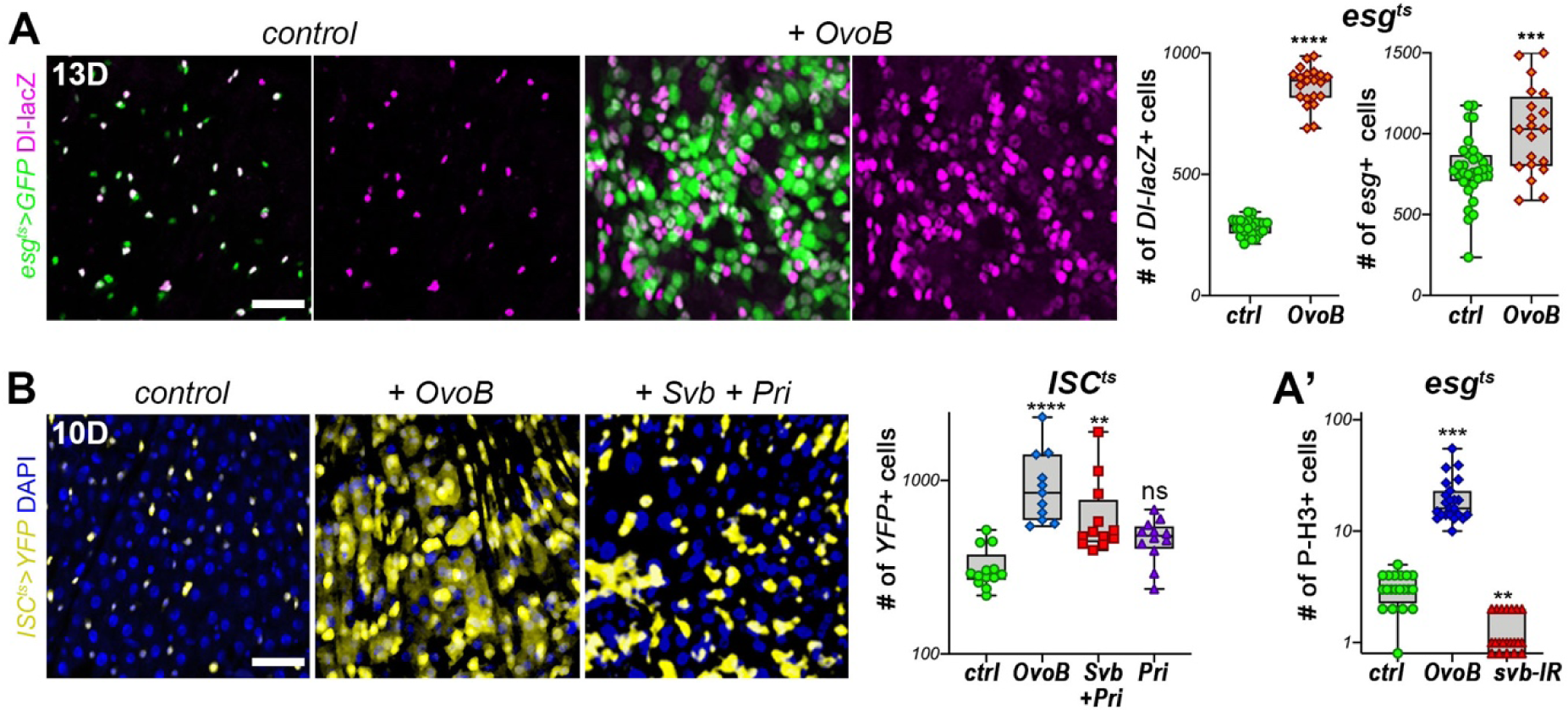
The Svb activator induces ISC proliferation. (A) Expression of *esg^ts^*>GFP (green) and *Dl-lacZ* (purple) in control or upon OvoB expression, and quantification of *Dl-lacZ* positive cells (ISCs) and GFP positive cells (ISC/EBs). (A’) Quantification of the mitotic marker P-H3 in control flies (*egs^tS^>GFP*), or expressing OvoB or *svb-RNAi* driven by *esg^ts^*. (B) Expression of YFP (yellow) driven by *ISC^ts^* in control, flies expressing *OvoB* or *Svb* plus *pri*, and quantification of YFP positive cells. Blue is DAPI. Scale bars represent 20µm. P values from Mann-Whitney tests (A’) and Kruskal-Wallis tests (B’) are ns: p>0.5, *: p<0.05, **: p<0.01, ***: p < 0.001, ****: p<0.0001. See also Figure S3.

These data thus show that the Svb^ACT^ form drives ISC proliferation and that Pri smORF peptides play a key role to regulate the processing of Svb in adult stem cells.

### Svb acts downstream of Wnt and EGFR mitogenic pathways in the adult midgut

*svb* was identified as a functional integrator of signaling pathways, including Wnt, EGFR, Hedgehog and Notch, mediating their roles in the terminal differentiation of embryonic epidermal cells (Payre, 2004). Since these pathways are well-established to control ISC maintenance, proliferation and differentiation (reviewed in (Buchon and Osman, 2015)), we investigated whether Svb could also mediate their activity in intestinal stem cells.

As previously reported (Lin et al., 2008), inhibition of Wnt signaling by expressing a dominant negative form of the nuclear effector TCF/Pangolin (TCF-DN) led to a marked decrease in the number of ISC/EBs (Figure 4A,A’). Co-expression of OvoB was not only sufficient to rescue the progenitor population, but could still induce ISC proliferation even when Wnt activity was compromised. Conversely, over-activation of Wnt signaling by expressing the Wingless ligand, or a constitutively active transcriptional regulator *Arm*, promotes ISC division (Lin et al., 2008). The Wnt-induced stem cell proliferation was further potentiated upon expression of OvoB and abrogated when *svb* function was inhibited (Figure 4A). Similarly, knocking-down EGFR through a dominant negative receptor (EGFR-DN) led to a loss of *esg+* cells, and OvoB was sufficient to bypass this depletion of stem cells (Figure 4B,B’). Activation of EGFR signaling (*esg^ts^F/O>RasV12*) induced progenitor proliferation, which was counteracted upon *svb* knockdown (Figure 4B). These results show that Svb is epistatic to, in other words likely acts downstream, Wnt and EGFR pathways to control adult stem cell maintenance and division.

**Figure 4:**
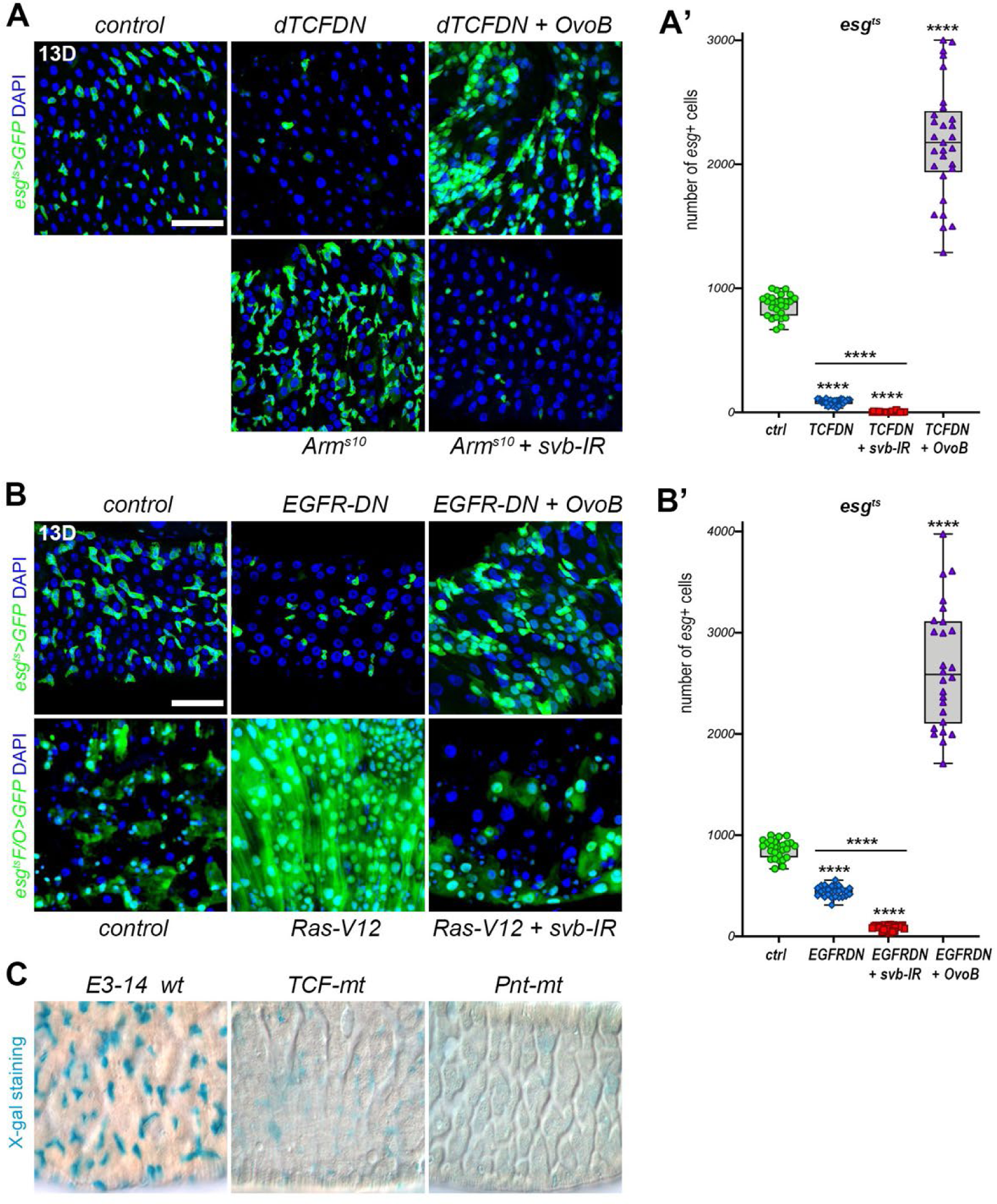
Svb acts downstream of mitogenic signaling pathways Wnt and EGFR in adult midgut. (A,A’) Images from control flies, or expressing TCFDN, TCFDN and OvoB, Arms10, Arms10 and svb-RNAi driven by esg*^ts^* and quantification of GFP-positive progenitors. (B, B’) Images from control *esg^ts^*>GFP flies or expressing EGFR-DN, EGFR-DN and OvoB and quantification of GFP-positive progenitors. Bottom panels show *egs^ts^F/O>GFP* (control), and expressing RasV12 alone or in combination with *svb*-RNAi. (C) X-gal staining of posterior midguts showing expression of *DmE3-14-LacZ*, or mutated version of the enhancer knocking out putative binding sites for Pointed or TCF. Scale bars represent 20µm. P values from Kruskal-Wallis tests ****: p<0.0001. See also Figure S4.

Therefore, *svb* expression integrates inputs from Wnt and EGFR pathways in both adult stem cells and embryonic epidermal cells (Payre et al., 1999); the underlying mechanisms remained, however, to be elucidated. To identify in an unbiased manner the transcription factors that control *svb* expression, we undertook a large-scale functional screening *in vivo*. We performed this screen in the embryo, where signaling pathways do not dramatically impinge on the presence/absence of targeted cells, as opposed to adult stem cells. Briefly, we selected the whole complement of transcription factors showing detectable expression at the time of *svb* expression in epidermal cells (227 candidates), and assayed consequences of their dsRNA-mediated depletion on the activity of the *E3-14 svb* enhancer (Figure S4A). This identified four factors the depletion of which alters *E3-14* expression, including TCF and Pointed, *i.e.*, the nuclear effectors of Wnt and EGFR signaling, respectively. Analysis of the *E3-14* sequence identified putative binding sites for TCF and Pointed (Figure S4B-C) and we generated transgenic flies carrying *E3-14-lacZ* reporters with mutations within either TCF (*E3-14-TCF-mt)* or Pointed (*E3-14-pnt-mt*) binding sites. When compared to the wild-type reporter in embryos, *E3-14-pnt-mt* displayed reduced expression, whereas *E3-14-TCF-mt* showed an ectopic pattern in epidermal cells (Figure S4D). To further assay whether this altered pattern affected *svb* function, we generated transgenic lines driving *svb* expression under the control of wild type *E3-14* or *E3-14-TCF-mt* enhancers and assayed their rescuing activity when introduced in *svb* mutant embryos. While *E3-14-svb* displayed a clear rescue of epidermal trichomes, the mutation of TCF binding sites almost completely abolished the function of *E3-14-TCF-mt* (Figure S4E). Accordingly, both *E3-14-TCF-mt* and *E3-14-Pnt-mt* displayed strongly decreased activity when assayed in adult stem cells (Figure 4C).

These data show that the medial *svb* enhancer integrates direct regulatory inputs from the Wnt and EGFR pathways, to control the behavior of intestinal stem cells.

### The Svb repressor promotes enterocyte differentiation

In addition to stem cells, we observed expression of Svb within differentiated enterocytes as deduced from complementary pieces of evidence. These data opened the possibility of a role of Svb in the differentiation of the intestinal stem cell lineage, which we investigated using a set of functional assays.

*In situ* hybridization revealed that *svb* mRNA is expressed in small doublet cells, *i.e.* ISC/EBs, but also in large-nuclei enterocytes all along the midgut epithelium (Figure 5A). This was further supported by the expression of a *svb*::*GFP* mini-gene rescue construct (Menoret et al., 2013) (Figure 5B), while no expression was detected in enteroendocrine cells. The *svb* proximal enhancer (*9CJ2*, see Figure 1A) showed a restricted expression in big nucleated cells (Figure 5C). We used the same screening assay in the embryo and found that *9CJ2* expression is reduced upon knocking down Pdm1/Nubbin (Figure S5), a TF well known for the differentiation of enterocytes (Korzelius et al., 2014; Tang et al., 2018). We identified two Pdm1 putative binding sites within the *9CJ2* enhancer (Figure S5B,C) and inactivated these sites by site-directed mutagenesis (*9CJ2-PDM-mt*). When assayed *in vivo*, the inactivation of Pdm1 binding sites prevents *9CJ2* expression in both embryonic epidermal cells and adult intestinal stem cells (Figures 5C and S5D). We therefore conclude that *svb* is also expressed in enterocytes under the control of the proximal enhancer, likely directly regulated by Pdm1.

**Figure 5:**
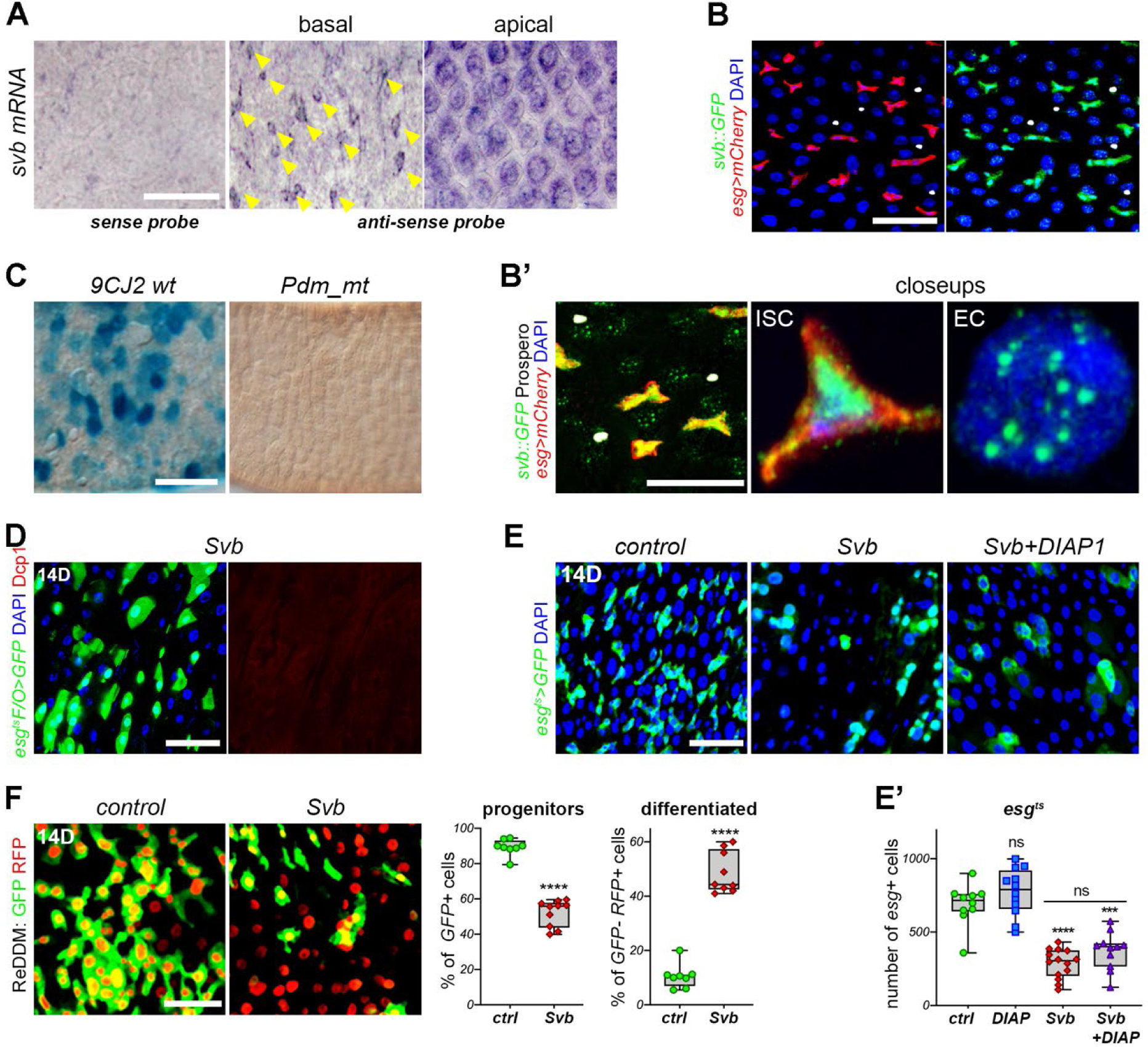
Svb repressor is expressed in enterocytes and promotes differentiation. (A) Expression of *svb* mRNA in the adult posterior midgut as revealed by *in situ* hybridization. Arrows indicate ISC/EBs. (B) Expression of a *svb* mini-gene rescue construct (Menoret et al., 2013) composed of *svb-cDNA* tagged by GFP (green) and driven by the medial and proximal *svb* enhancers. Red is mCherry and white Prospero. (C) Expression of wild type (*9CJ2*) and a mutated version disrupting the putative Pdm1 binding site (*9CJ2-mut-PDM1*) *svb* proximal enhancer. (D) Images of *esg^ts^-F/O* clones expressing Svb and stained for DCP1(red), clones are visualized by GFP (green). (E,E’) Control (*esg^ts^>GFP*) and midguts expressing Svb alone or combined to DIAP1 driven by *esg^ts^* (E) and quantification (E’). (F) Expression of ReDDM in control flies or expressing Svb and quantification of progenitors and differentiated cells. In all panels, blue is DAPI and scale bars represent 20µm. P values from Mann-Whitney tests (F’) and Kruskal-Wallis tests (E’) are ns: p>0.5, ***: p < 0.001, ****: p<0.0001. See also Figure S5.

The switch in Svb transcriptional activity is associated with a striking change in its intra-nuclear distribution: whereas Svb^ACT^ diffusely distributes within the nucleoplasm, Svb^REP^ accumulates in dense foci (Kondo et al., 2010; Zanet et al., 2015). Interestingly, Svb staining is diffuse in *esg+* cells (which express *pri*), while it accumulates in intra-nuclear foci in enterocytes (Figures 2B and 5B’). This differential pattern of nuclear distribution thus suggested that, unlike ISCs, Svb might act as a repressor within ECs where it could play a role in their survival and/or differentiation. To test this hypothesis, we overexpressed the full size Svb protein (Svb^REP^) in intestinal progenitors. The number of ISC/EB was reduced and remaining *esg+* cells displayed aberrant morphology, with enlarged nuclei reminiscent of polyploid ECs (Figure 5E). Similar phenotypes were observed when using the *ISC^ts^* driver. Importantly, the loss of progenitors observed upon Svb^REP^ expression was not rescued by concomitant expression of *DIAP1* (Figure 5E,E’). *esg^ts^F/O* clones expressing Svb were composed of single cells displaying large nuclei, and being negative for DCP1 apoptotic staining (Figure 5D). These data thus suggested that Svb induces ISC differentiation into enterocytes rather than cell death, a hypothesis we further tested using ReDDM lineage tracing (Antonello et al., 2015). The loss of progenitors upon Svb^REP^ expression was accompanied by an increase in differentiated cells (Figure 5F-F’) showing that the Svb repressor induces progenitor differentiation towards the enterocyte fate.

To further investigate Svb function within enterocytes, we expressed *svb-*RNAi using the EC-specific driver *MyoIA^ts^ (MyoIA-Gal4, UAS-GFP, tubulin-Gal80^ts^)*. We observed that *svb* knockdown induced EC apoptosis, as shown by elevated DCP1 staining (Figure S6A). We next further assayed whether Svb acted as a repressor in ECs, through driving Ovo/Svb isoforms in ECs with *MyoIA^ts^*. Intestines expressing Svb^REP^ did not show any detectable homeostatic or structural changes. In contrast, OvoB induced a loss of GFP+ ECs, with an increase in EEs (Figure 6A). In addition, the apical microvilli that features mature ECs was altered following *svb* knockdown in enterocytes (Figure 6B,C). Furthermore, ectopic expression of Svb^ACT^ also disrupted brush border differentiation, whereas we did not detect defects upon Svb^REP^ expression.

**Figure 6:**
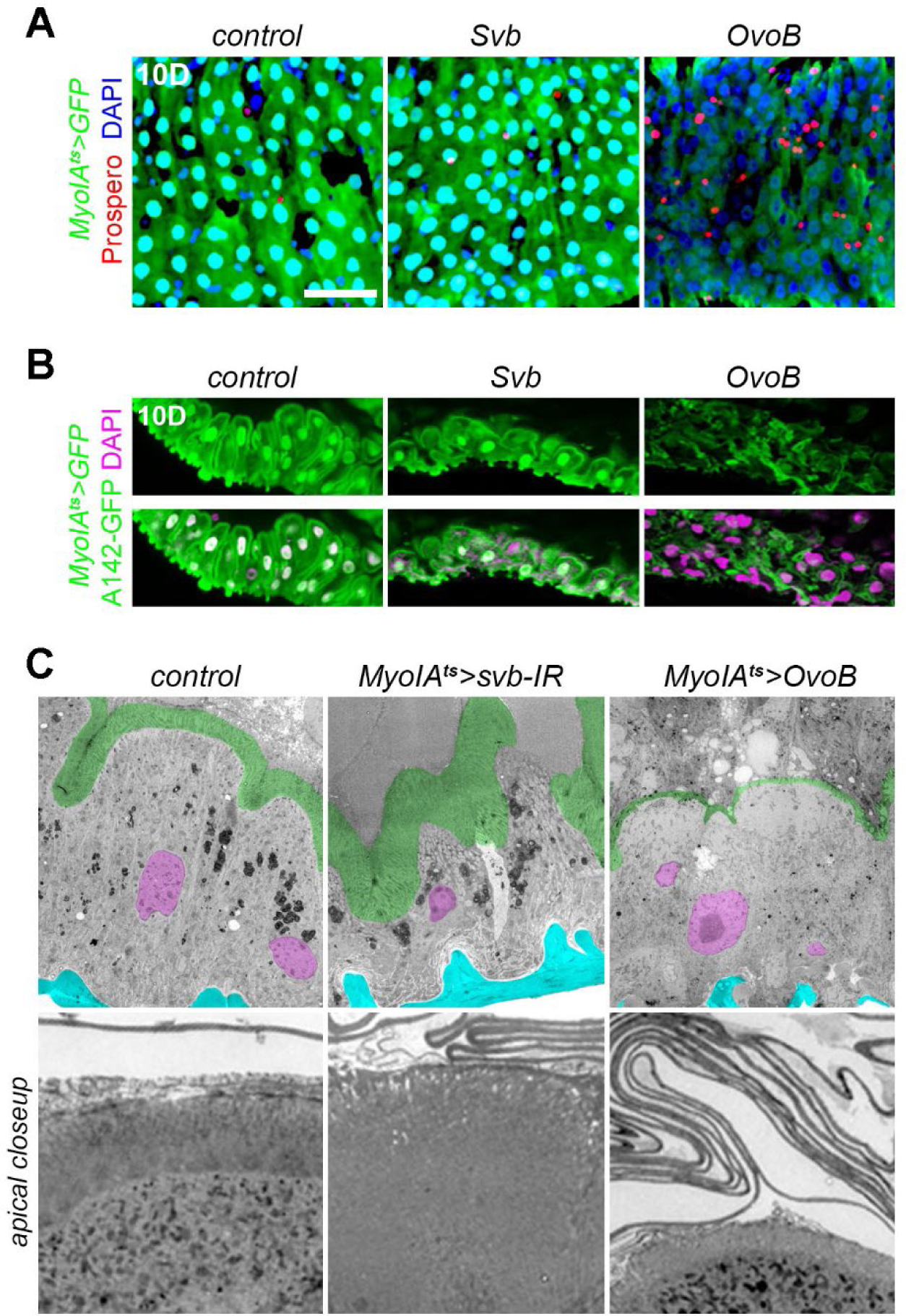
The Svb repressor is required to maintain proper enterocyte differentiation. (A) Control midgut or midgut expressing *Svb* or *OvoB* in EC. EC are visualized by GFP. Red is Prospero and blue is DAPI. Scale bar represents 20µm. (B) Images from control midgut or midgut expressing Svb or OvoB. ECs are visualized by GFP and their brush border apical organization underlined by the expression of *A142::GFP*. (C) Electron micrographs of control (*MyoIA^ts^>GFP*), *MyoIA^ts^>svb-RNAi* and *MyoIA^ts^>OvoB*. Either the depletion of *svb* or the ectopic expression of *OvoB* leads to strong defects in enterocyte ultrastructure, including alteration of the brush border microvilli (highlighted in green). Nuclei are pseudo-colored in purple and the basement membrane in cyan.

All together, these findings indicate that the Svb repressor is required to maintain proper differentiation/maturation state of ECs in the adult midgut.

### Svb allows the growth of genetically-induced ISC tumors

A main determinant of intestinal stem cell differentiation is the activation of the Notch pathway (Ohlstein and Spradling, 2007). We hypothesized that Svb may synergize with Notch to direct EC differentiation.

As previously reported (Ohlstein and Spradling, 2007), inhibition of Notch induces tumor-like expansion of ISCs (Figure 7A). Concomitant *svb* knockdown strongly reduced the expansion of Notch-induced tumors. Strikingly, Svb^REP^ was sufficient to suppress Notch tumors and induced EC differentiation (Figure 7A). Conversely, Notch activation by expressing the nuclear Notch intracellular domain (NICD) caused an acute decrease in progenitors, likely differentiating in ECs (Figure S7A). The co-expression of NICD and OvoB restored a wild type number of progenitors, whereas co-expressing Svb^REP^ further enhanced the NICD phenotype (Figure S7A). Furthermore, we found antagonistic effects of Svb^ACT^ *versus* Svb^REP^ on the phenotype resulting from the expression of a dominant negative form of Delta, the Notch ligand in ISCs (Figure S7B).

**Figure 7:**
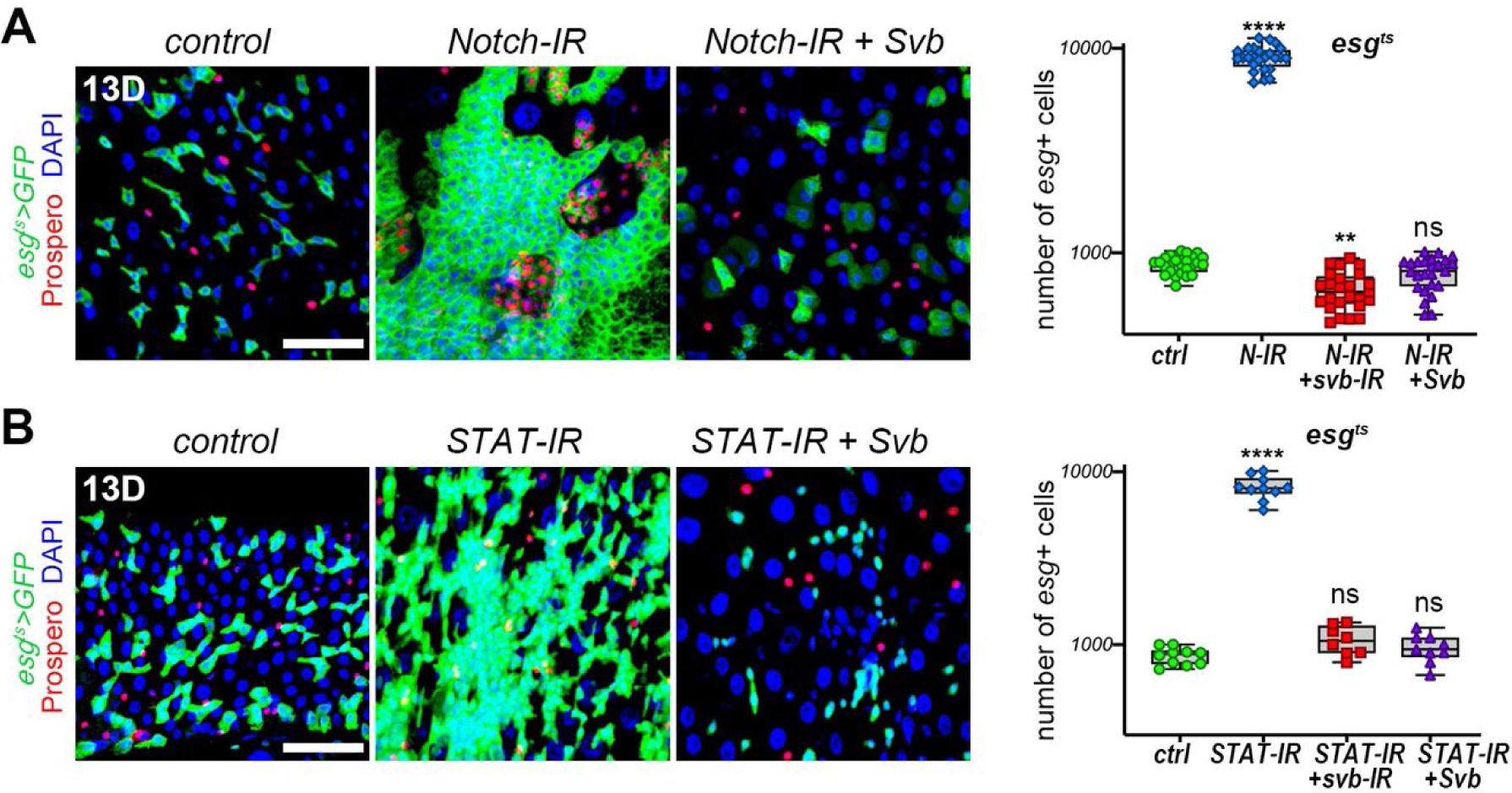
The Svb repressor suppresses genetically-induced hyperplasia in the gut. (A) *Esg^ts^*-driven expression of *Notch*-RNAi alone, or combined to *svb*-RNAi or to Svb and quantification of GFP positive cells. (B) Effect of *esg^ts^* expression of *STAT*-RNAi alone, or combined to *svb*-RNAi or to S and quantification of GFP positive cells. In all panels, GFP is in green, anti-Prospero staining in red and DAPI in blue; scale bars represent 20µm. P values from Kruskal-Wallis tests are ns: p>0.5, **: p < 0.01, ****: p<0.0001. See also Figure S7.

We next tested whether the Svb repressor was sufficient to counteract the deregulation of additional signaling pathways that induce midgut tumors. Indeed, we found that Svb^REP^ expression inhibited the expansion of progenitors induced by inactivation of the STAT transcription factor, or the overexpression of Wg, or Arm^s10^, in *esg+* cells (Figures 7B and S7C,D). In addition, the depletion of *svb* in those tumors was sufficient to block their expansion in the midgut (Figure 7B).

We conclude that *svb* is required for the growth of genetically-induced ISC tumors and that the Svb repressor overrides different signaling pathways to promote enterocyte differentiation.

## DISCUSSION

Our data show that the OvoL/Shavenbaby transcription factor is a key regulator of adult intestinal stem cells and their progeny. In stem cells, Svb is post-translationally processed into an activator required for progenitor maintenance. Svb^ACT^ is sufficient to promote stem cell division, thereby mediating the proliferative activity of EGFR and Wnt signaling. Svb is also expressed in enterocytes, where the unprocessed Svb repressor promotes differentiation. The balance between Svb^ACT^ and Svb^REP^ is gated by Pri smORF peptides, which allow proteasome-mediated conversion of Svb transcriptional activity in response to Ecdysone signaling (Chanut-Delalande et al., 2014; Kondo et al., 2010; Zanet et al., 2015). Our results further suggest that the Svb transcriptional switch is a conserved mechanism involved in the control of various adult stem cells.

### *svb* integrates multiple regulatory cues for the homeostasis of adult stem cells

*svb* expression is driven by a large array of enhancers (Stern and Frankel, 2013), which collectively define at single cell resolution the pattern of epidermal differentiation (Frankel et al., 2011; McGregor et al., 2007; Preger-Ben Noon et al., 2016; Sucena et al., 2003). Although *svb* enhancers were delineated for their embryonic activity, they harbor pleiotropic functions across the *Drosophila* life-cycle (Bohere et al., 2018; Preger-Ben Noon et al., 2018). At least three enhancers drive *svb* expression within the lineage of adult intestinal stem cells. In stem cells and progenitors, *svb* enhancers integrate regulatory outputs of Wnt and EGFR signaling to direct *svb* expression. Unbiased large-scale screening further indicates that the nuclear mediators of Wnt (TCF) and EGFR (Pointed) directly regulate the activity of a *svb* stem cell enhancer. In addition, PDM1, a key TF for the enterocyte fate (Korzelius et al., 2014), regulates a distinct enhancer driving *svb* expression within differentiated enterocytes. Hence, the regulatory logic of *svb* expression is reused in the adult intestinal lineage, involving separate enhancers and regulators between stem cells and enterocytes.

### Svb^ACT^ is required for the maintenance and proliferation of stem cells

A main aspect of Svb regulation relies on post-translational control of its transcriptional activity (Bohere et al., 2018; Chanut-Delalande et al., 2014; Kondo et al., 2010; Menoret et al., 2013; Ray et al., 2019; Zanet et al., 2015). Svb processing into an activator is indispensable for the maintenance of stem cells, which otherwise undergo apoptosis. Both intestinal and renal stem cells are particularly resistant to apoptotic cell death (Ma et al., 2016). In the latter, we recently reported that Svb physically interacts with Yorkie (a.k.a. YAP/TAZ in mammals), the nuclear effector of Hippo signaling (Staley and Irvine, 2012), to directly activate DIAP1 expression (Bohere et al., 2018). Although renal stem cells are mostly quiescent (Bohere et al., 2018; Xu et al., 2018), intestinal stem cells self-renew under homeostatic conditions and proliferate in response to various challenges (Staley and Irvine, 2012). Consistently, Svb^ACT^ is required, and to a certain extent sufficient, to promote stem cell proliferation. Knocking out *svb* function, or maturation, prevents normal stem cell maintenance and suppresses their expansion induced by mitogenic signals (Wnt, EGFR, JAK/STAT…). Reciprocally, Svb^ACT^ can bypass the repressive effects of altered signaling and triggers stem cell expansion. These data support a model whereby the Svb activator plays a key role in mediating the activity of signaling pathways for the control of adult stem cell division.

### Svb^REP^ in the differentiation of enterocytes

Our results also show the expression and function of Shavenbaby in enterocytes, but not in enteroendocrine cells, consistent with an early separation between EC and EE lineages (Guo and Ohlstein, 2015; Yin and Xi, 2018; Zeng and Hou, 2015). In contrast to stem cells, Svb acts as a repressor within enterocytes where it is required for their functional organization, including proper differentiation of brush border microvilli. Strikingly, Svb^REP^ is sufficient to induce differentiation toward the enterocyte fate, including in conditions otherwise leading to stem cell expansion. This is the case for deregulated signaling, when Svb^REP^ can force differentiation of tumor-like stem cells. These findings therefore show that Svb^ACT^ and Svb^REP^ exert antagonistic function within the adult intestinal lineage, Svb^ACT^ promoting stem cell proliferation and progenitor survival, while Svb^REP^ later acts to promote and maintain enterocyte differentiation.

The switch between Svb^REP^ and Svb^ACT^ is triggered by Pri small peptides throughout the fly life-cycle (Bohere et al., 2018; Chanut-Delalande et al., 2014; Kondo et al., 2010; Zanet et al., 2015) and across arthropods (Ray et al., 2019; Savard et al., 2006). *pri* expression is manifest in progenitors but absent from differentiated enterocytes, and Pri peptides are required, together with their target Ubr3 ubiquitin ligase, for stem cell maintenance. A key role of Pri is to mediate systemic Ecdysone signaling for developmental timing (Chanut-Delalande et al., 2014). The Ecdysone receptor binds to *pri* cis-regulatory enhancers and trigger *pri* expression upon elevation of Ecdysone titer (Chanut-Delalande et al., 2014). Although Ecdysone influences intestine stem cells (Reiff et al., 2015), the role of Ecdysone in midgut homeostasis remains to be fully elucidated (Miguel-Aliaga et al., 2018). We find that disrupting Ecdysone signaling affects the behavior of intestinal stem cells, decreasing proliferation and promoting differentiation, *i.e.*, as observed upon inhibition of Svb processing. The expression of *pri*, or of constitutive Svb^ACT^, can compensate compromised Ecdysone signaling, showing that the regulation of Svb activity is a key target of hormonal control in intestinal stem cells.

### OvoL/Svb transcriptional switch for stem cell control across animals

Mounting evidence suggests a broad role of the Svb transcriptional switch in stemness. In flies, Shavenbaby refers to the somatic protein (Delon et al., 2003; Mevel-Ninio et al., 1995), while alternate promoters produce germinal isoforms called OvoA and OvoB, acting as a repressor and activator, respectively (Andrews et al., 2000; Mevel-Ninio et al., 1991; Mevel-Ninio et al., 1995). OvoB is required for the maintenance of germ cells (Delon et al., 2003; Hayashi et al., 2017; Mevel-Ninio et al., 1995), while OvoA later acts for proper differentiation (Andrews et al., 2000; Hayashi et al., 2017). Precocious expression of OvoA leads to germ line loss (Andrews et al., 1998; Mevel-Ninio et al., 1996) and other *ovo* mutations cause ovarian tumors. Although relying on different mechanisms between soma (post-translational processing) and germline (alternative promoters), the REP-to-ACT switch appears central to the function of Ovo/Svb factors in *Drosophila* stem cells. It will be interesting to elucidate whether a putative switch in transcriptional properties also underlies OvoL function in stem cells across animals.

OvoL factors are associated with many cancers, in particular those of epithelial origin (Roca et al., 2013; Roca et al., 2014). In flies, *svb* knockdown is sufficient to block Notch- and JAK/STAT-induced tumor formation and growth. Moreover, there are opposing effects of Svb^ACT^ *vs* Svb^REP^, which promotes or suppresses tumors, respectively. Interestingly, OvoL2 isoforms display strikingly different effects within tumors, in which only the repressor suppresses tumor development (Watanabe et al., 2014). Wnt signaling plays a key role in tumors, including in cancer stem cells (Zhan et al., 2017). We show that Svb is a target of Wnt and EGFR pathways in the fly gut, mediating their action on stem cell proliferation. These data show the role of antagonistic OvoL/Svb factors in the control of normal and tumor stem cells.

OvoL/Svb have been proposed to act as epithelial stabilization factors, counteracting epithelial to mesenchyme transition (EMT) (Lee et al., 2014; Li and Yang, 2014). EMT is induced by a core of transcription factors, including Snail, Slug and Zeb1,2 in mammals, and their *Drosophila* homologs (Escargot, ZFh1,2) prevent differentiation of intestinal stem cells (Antonello et al., 2015; Korzelius et al., 2014; Loza-Coll et al., 2014). According to the model of an antagonism between EMT factors and OvoL/Svb, the Svb^REP^ isoform promotes differentiation. However, our results draw a more complex picture, where Svb^ACT^ also acts together with Esg, ZFh1,2 for the maintenance of stem cells. EMT is not an all-or-none process and instead progresses through a series of reversible intermediate states between the epithelial (E) and mesenchyme (M) phenotypes (Nieto et al., 2016). Such hybrid E/M phenotypes are hallmarks of normal and cancer stem cells, and relative doses of EMT factors and OvoL/Svb may provide a tunable window of stemness (Jolly et al., 2015).

Taken together, these data therefore support a conserved role of OvoL/Svb transcription factor in the control of stem cell behavior, in both normal and tumorous conditions.

## ACKNOWLEDGMENTS

We thank the Bloomington Drosophila Stock Center, Kyoto Drosophila Genomics and Genetic Resources, Vienna Drosophila Resource Center, Flybase, Developmental Studies Hybridoma Bank, and A. Bardin, H. Bellen, J.P. Couso, A. Debec, N. Frankel, M. Dominguez, B. Edgar, D. Stern, and N. Tapon for providing fly stocks and reagents. We are grateful to M. Bou Sleiman, Z. Zongzhao and members of the FP lab for discussions and critical reading of the manuscript, J. Favier and A. Destenabes for help with transgenics, as well as S. Bosch for imaging. SAH was supported by Fondation pour la Recherche Médicale and Communauté des Communes de Dannieh-Université Libanaise. This work was supported by Agence Universitaire de la Francophonie (PCSI AUF-BMO), École Doctorale des Sciences et de Technologie-Université Libanaise, The Federation of European Biochemical Societies, American University of Beirut URB, Agence Nationale de la Recherche (ANR, ChronoNet) and Fondation pour la Recherche Médicale (DEQ20170336739). The funders had no role in study design, data collection and interpretation, or the decision to submit the work for publication.

## AUTHOR CONTRIBUTIONS

The project was initially setup by DO, SP and FP. FP and DO conceptualized and supervised the work. SAH, FP and DO designed experiments and wrote the manuscript; all authors commented and approved the final version. SAH, AA, JPB, JB, and DO performed experiments. SAH, AA, SP, FP and DO analyzed data. BR helped with imaging and image analysis, and ZK and BL shared key resources and materials.

