## Supplemental information for "Shavenbaby protein isoforms orchestrate the self-renewal *versus* differentiation of *Drosophila* intestinal stem cells"

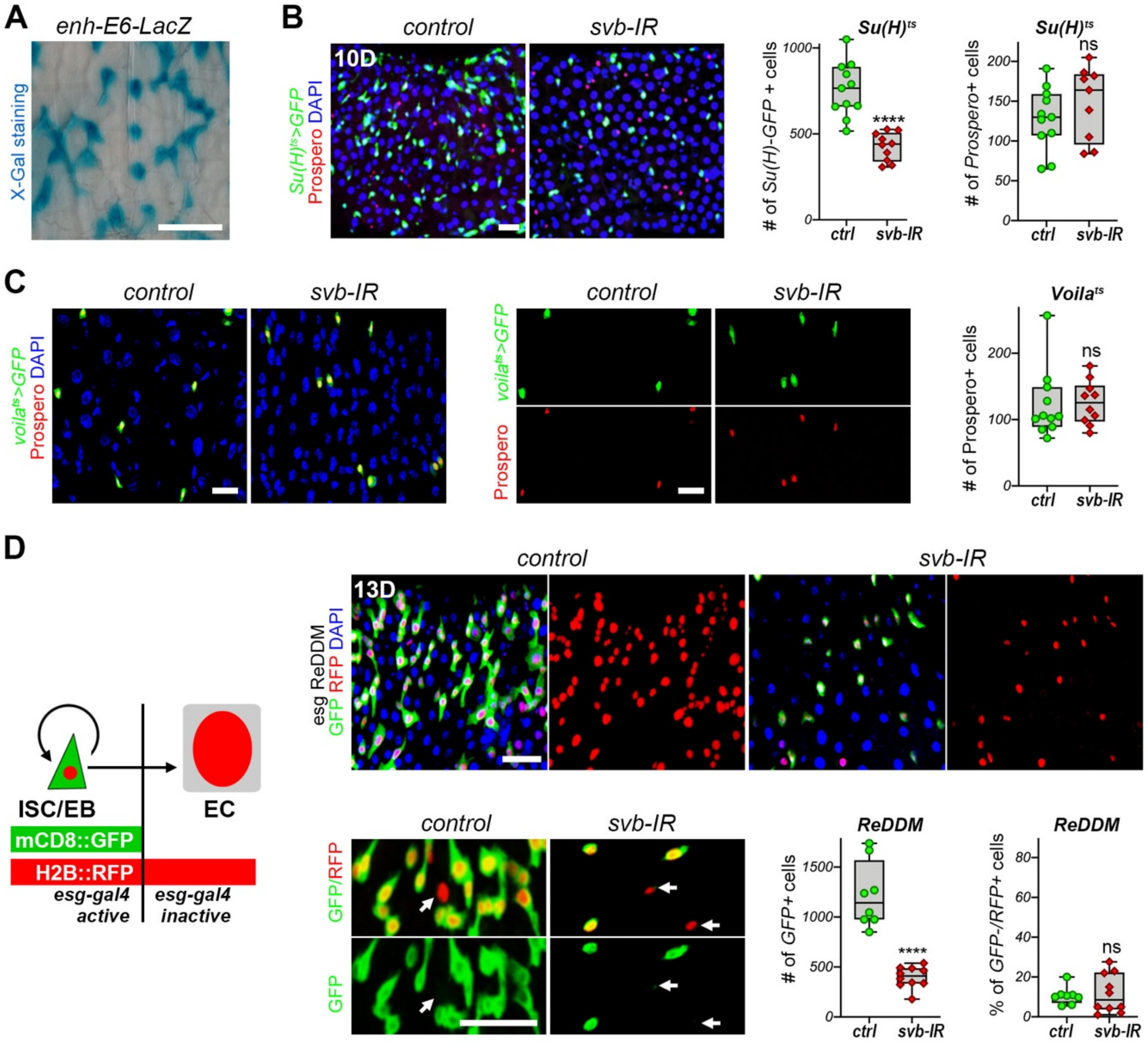


### Supplementary Figure 1

(A) Expression of the *E6 svb* enhancer, as reported by X-gal staining (cyan) of the *E6-LacZ* reporter line.
(B) *svb* knockdown in enteroblasts, driven by *Su(H)*$^{Ts}$. *Su(H)-GBE-GFP* cells are in green and anti-Prospero in red.
(C) Knocking-down *svb* function in enteroendocrine cells driven by *Voila*$^{Ts}$ does not affect their number. GFP positive cells are in green and anti-Prospero in red.
(D) Schematic representation of the ReDDM lineage tracing system (Antonello et al., 2015), in which *esg*$^{Ts}$ drives expression of both *mCD8::GFP* and *H2B::RFP*. *Esg+* cells are labelled by GFP (green) and RFP (red), while differentiated progeny only maintains the very stable H2B::RFP. *svb* knockdown results in a strong decrease on *esg*+ cells (GFP+) upon two weeks of treatment, while it does not affect the ratio between progenitors (GFP+) and differentiated progeny (RFP+/GFP-).
In all panels, blue is DAPI. P values from Mann-Whitney tests are: ns>0,05, ****<0,0001.

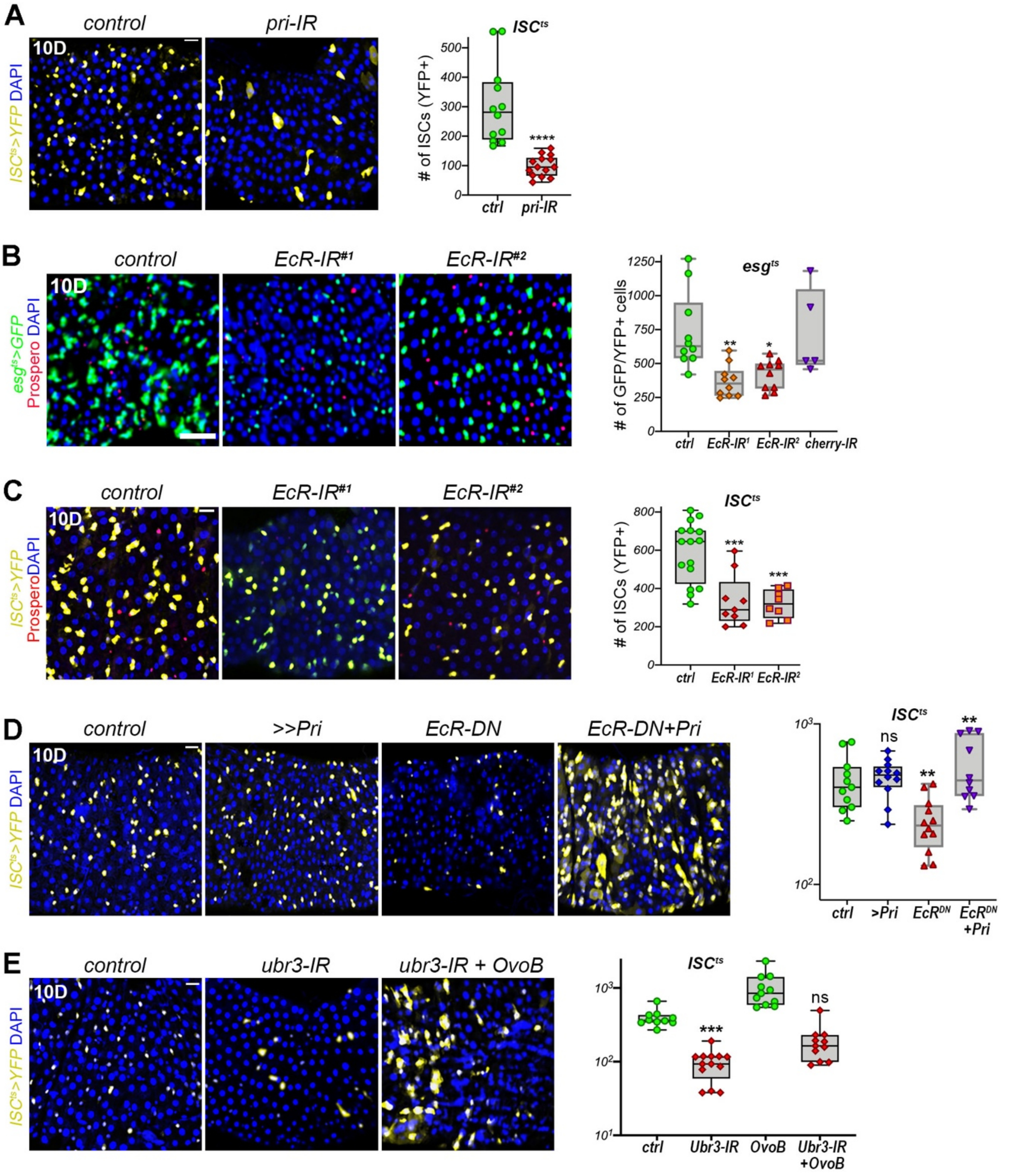


### Supplementary Figure 2

(A) Knockdown of *pri* by *ISC$^{Ts}$-Gal4* leads to the loss of YFP+ stem cell (yellow).
(B) Depletion of the Ecdysone receptor (*EcR*) using two RNAi lines driven by *esg$^{TS}$-Gal4* reduces the pool of intestinal progenitors (green). An RNAi line targeting mcherry was used as a negative control.
(C) *EcR* knockdown driven by *ISC$^{TS}$-Gal4* causes the loss of *YFP+* stem cells (yellow).
(D) Stem cell loss upon *EcR-DN* expression driven by *ISC$^{TS}$-Gal4* is compensated by re-expression of *pri.* For easier visualization, the graph is shown using a log (10) y scale.
(E) Depletion of *Ubr3* driven by *ISC$^{TS}$-Gal4* reduces the number of YFP+ stem cells (yellow) and can be compensated by concomitant expression of the constitute activator *OvoB*. For easier visualization, the graph is shown using a log (10) y scale.
All panels display phenotypes following 10 days of treatment, with DAPI in blue and Anti-Prospero staining in red. P values from Mann-Whitney tests (A) or Kruskal-Wallis ANOVA are: ns>0.05; *<0.05; **<0.01; ***<0.001; ****<0.0001.

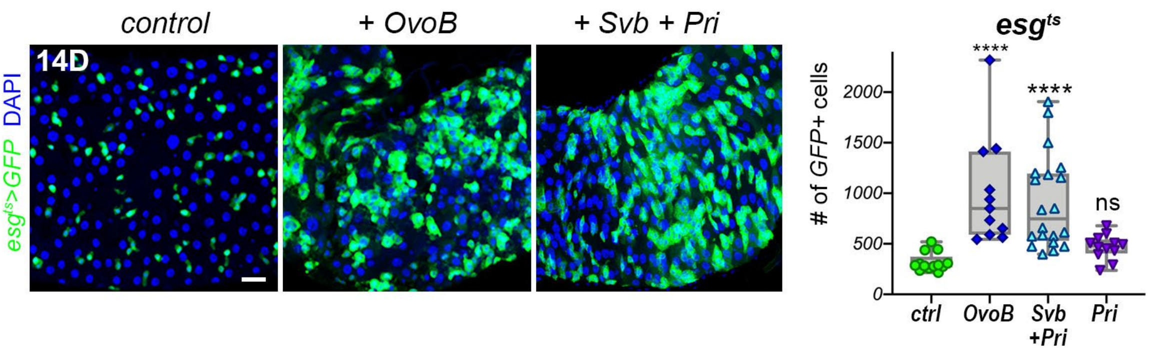


### Supplementary Figure 3

Expression of *OvoB*, or *Svb* plus *Pri*, driven by *esg$^{ts}$-Gal4* induces over-proliferation of GFP+ (green) progenitors and quantification of GFP+ cells.Blue is DAPI. P values from Kruskal-Wallis ANOVA are: ns>0.05; ****<0.0001.

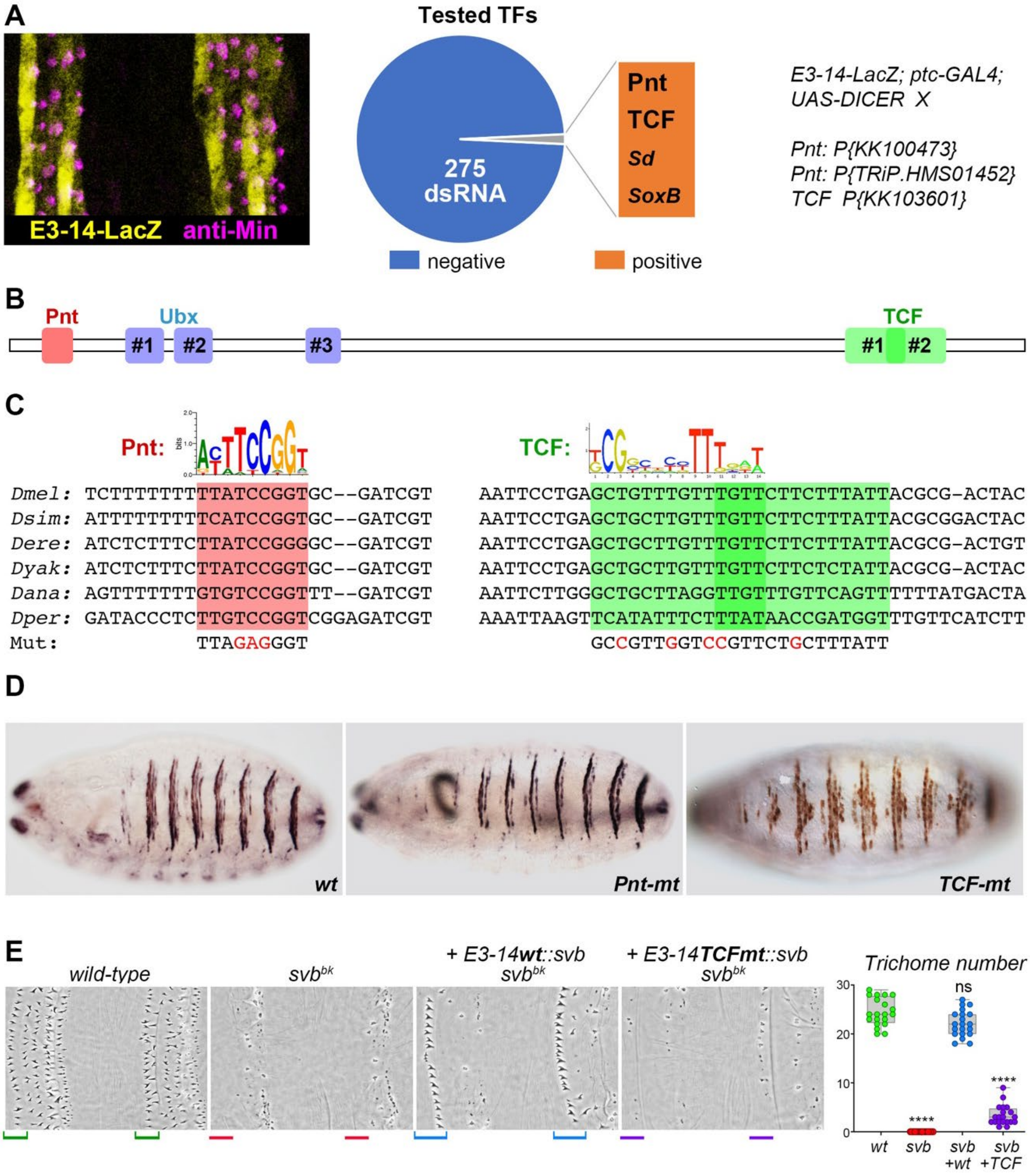


### Supplementary Figure 4

(A) The *E3_14 svb* enhancer is expressed in ventral epidermal trichome cells, with strong expression in the anterior-most (at left) row of trichomes within each segment. Lac-Z staining is in yellow, the Miniature protein (purple) highlights growing trichomes. Schematic view of functional screening for 265 TFs using *ptc-GAL4* to drive corresponding RNAi. Knockdown of four candidate TFs alter the expression pattern of the *E3_14 svb* enhancer.

(B) Drawing of the *E3_14 svb* enhancer, with position of binding sites for Ubx (Crocker et al., 2015), Pointed (Pnt) and TCF.

(C) Evolutionary conservation of DNA sequence encompassing Pnt (red) and TCF (green) binding sites within the *E3_14 svb* enhancer. Nucleotides in red represent point mutations that have been introduced in *E3_14 svb* enhancer to disrupt either Pnt or TCF binding sites.

(D) Consequences of knocking out Pnt or TCF binding sites on the expression of the *E3_14 svb* enhancer. Pictures show ventral view of stage 15 embryos.

(E) Trichome rescue assays showing the influence of TCF binding site knockout. Picture show cuticle preparations of wild type, and *svb* mutant embryos, focusing on ventral of the third and fourth abdominal segments. *Svb* mutants display strong reduction in the number of trichomes, remaining trichomes are highly abnormal. Consistent with its expression pattern, the *E3_14 svb* enhancer driving *svb* cDNA (*E3-14wt::svb*) is able of rescuing trichomes in a *svb* mutant background, with striking rescue of the anterior-most trichome row (bracketed). Knocking out TCF binding sites (*E3-14TCFmt::svb*) strongly decreases trichome rescuing ability of the *E3_14 svb* enhancer. P values from Kruskal-Wallis ANOVA are: ns>0.9; ****<0.0001.

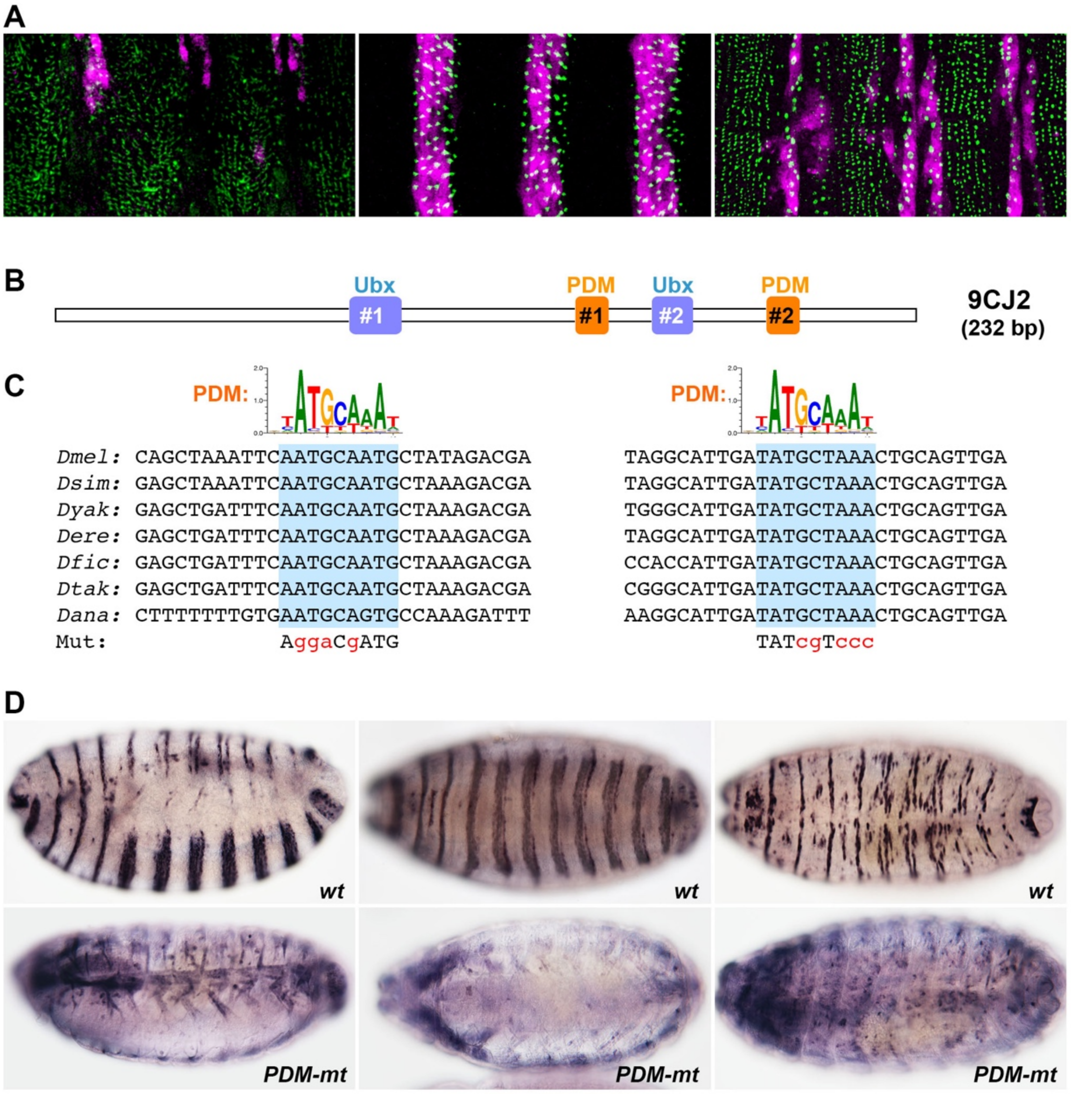


### Supplementary Figure 5

(A) The *9CJ2 svb* enhancer (purple) is expressed in most ventral epidermal trichome cells and in two stripes of trichome cells in the dorsal region of each abdominal segment. The Dusky-like protein (green) highlights growing trichomes.
(B) Drawing of the *9CJ2 svb* enhancer, with position of binding sites for Ubx (Crocker et al., 2015) and Nubbin/PDM-1 (PDM).
(C) Evolutionary conservation of DNA sequence encompassing each of the two PDM (orange) binding sites within the *9CJ2 svb* enhancer. Nucleotides in red represent mutations that have been introduced in *9CJ2 svb* enhancer to disrupt PDM binding sites.
(D) Consequences of knocking down PDM binding sites on the expression of the *9CJ2 svb* enhancer. Pictures show lateral (left panels), ventral (middle panels) and dorsal (right) views of stage 15 embryos.

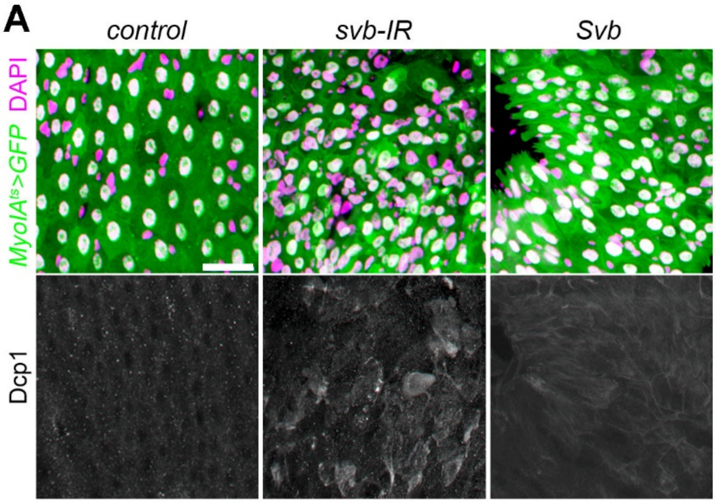


### Supplementary Figure 6

(A) *Myo1A*$^{ts}$ expression of GFP alone (control) or in combination with -RNAi, or Svb. GFP is in green, DCP1 in red (or white in the lower panels) and DAPI is purple. Scale bar is 20µm.

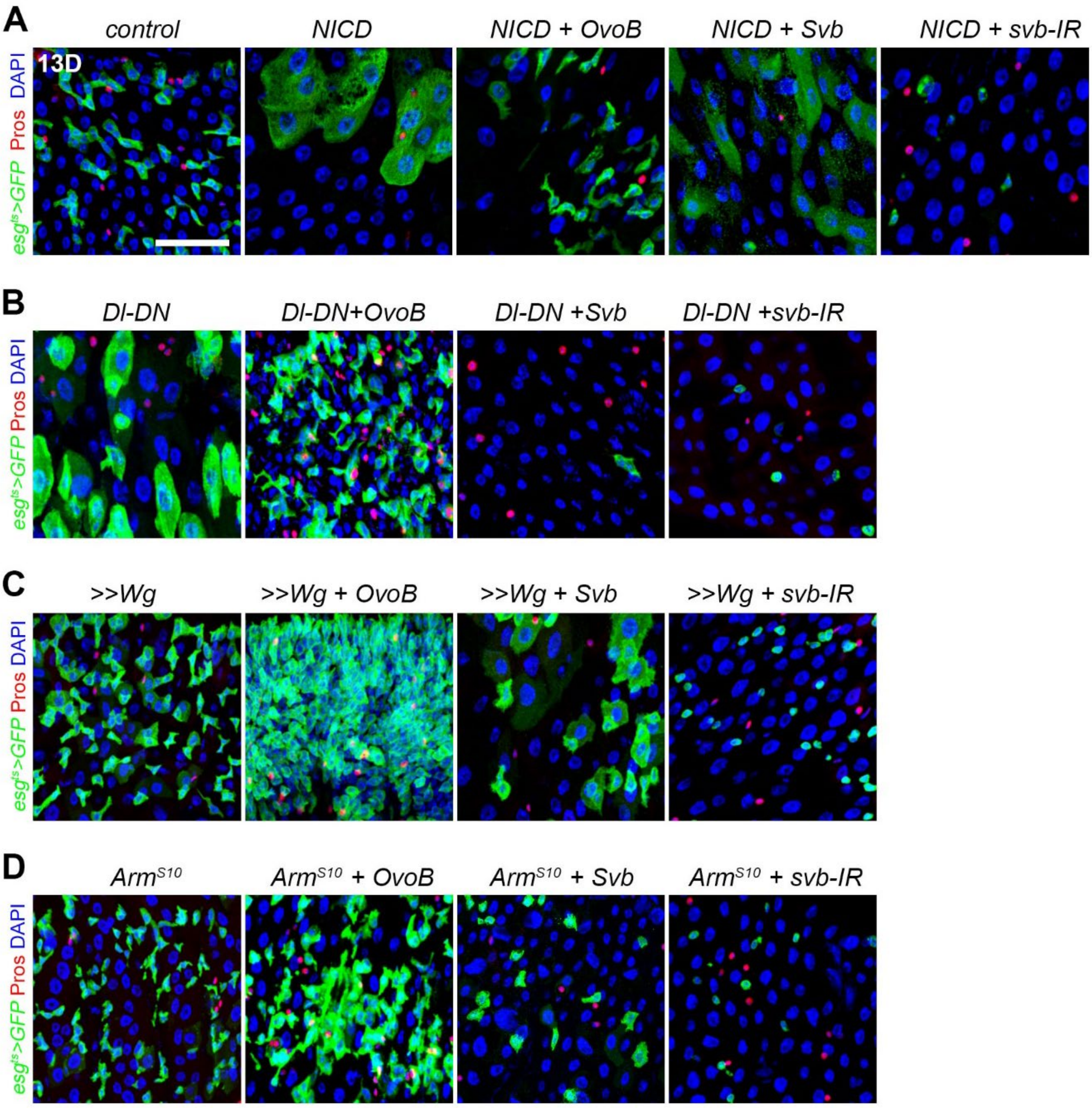


### Supplementary Figure 7

(A) *Esg*$^{TS}$*-Gal4* driven expression of Notch Intra Cellular Domain (NICD, which mimics activation of the Notch pathway) alone, or in combination with OvoB of Svb$^{REP}$ or *svb-IR*.
(B) Expression of a dominant negative form of the Delta ligand (Dl-DN) alone, or in combination with OvoB, Svb, or *svb-RNAi,* driven by *esg*$^{TS}$*-Gal4.*
(C,D) *Svb*$^{REP}$ counteracts progenitor over-proliferation upon activation of the *Wnt* pathway qnd impose differentiation. *Esg*$^{TS}$*-Gal4* driven expression of Wg or Arm$^{S10}$, with or without concomitant expression of OvoB, Svb or *svb-RNAi.*
In all panels GFP+ cells are in green, anti-Prospero staining in red and DAPI in blue. Scale bar is 20µm.
